# TELSAM polymers accelerate crystallization of fused target proteins by stabilizing minimal crystal contacts and in the absence of direct inter-TELSAM contacts

**DOI:** 10.1101/2021.07.23.453529

**Authors:** Supeshala D. Sarath Nawarathnage, Sara Soleimani, Moriah H. Mathis, Braydan D. Bezzant, Diana T Ramírez, Parag Gajjar, Derick Bunn, Cameron Stewart, Tobin Smith, Maria J Pedroza Romo, Seth Brown, Tzanko Doukov, James D. Moody

## Abstract

We extend investigation into the usefulness of genetic fusion to TELSAM polymers as an effective protein crystallization strategy. We tested various numbers of the target protein fused per turn of the TELSAM helical polymer and various TELSAM–target connection strategies. We provide definitive evidence that: 1. A TELSAM–target protein fusion can crystallize more rapidly than the same target protein alone, 2. TELSAM–target protein fusions can form well-ordered, diffracting crystals using either flexible or rigid TELSAM–target linkers, 3. Well-ordered crystals can be obtained when either 2 or 6 copies of the target protein are presented per turn of the TELSAM helical polymer, 4. The TELSAM polymers themselves need not directly contact one another in the crystal lattice, and 5. Fusion to TELSAM polymer confers immense avidity to stabilize exquisitely weak inter-target protein crystal contacts. We report features of TELSAM-target protein crystals and outline future work needed to define the requirements for reliably obtaining optimal crystals of TELSAM–target protein fusions.

## Introduction

Atomic-resolution protein structures are essential for structure-function studies, structure-based drug design, and biomedical protein engineering. X-ray crystallography remains an important technique to determine atomic-level protein structure, especially of proteins too small for single-particle cryo-electron microscopy. Protein crystals are also needed for micro-electron diffraction [1] and time-resolved diffraction using X-ray free-electron lasers [2]. Current protein crystallization methods are successful for only about 10% of all known proteins [3], and constitute a lengthy, laborious, and expensive process [4]. Lack of high-resolution structures hampers the structure-function studies of many proteins. There is a critical need for new protein crystallization methods that require less labor, time, and resources and that can induce the crystallization of a wider range of proteins.

A pH-sensitive mutant of the polymer-forming sterile alpha motif domain (SAM) of human Translocation ETS Leukemia (TEL) protein (TELSAM) was previously engineered [5]. It was then shown that genetic fusion of a panel of target proteins of interest to this TELSAM protein polymer could consistently achieve their crystallization [6, 7]. While many of the resulting crystals were too disordered to permit structure determination, we posit that continued investigation into the requirements for obtaining well-ordered crystals of TELSAM-target protein fusions is warranted for the following reasons: 1. The long TELSAM polymers are expected to confer immense avidity to any weak crystal contacts made by fused target proteins. 2. The regular spacing of target proteins along the 6-fold helical TELSAM polymer is expected to pre-program much of the symmetry of the resulting crystal lattice. 3. Fusion to TELSAM forces target proteins to participate in the resulting crystal lattice, ordering the target proteins and allowing them to be resolved in the resulting electron density maps. 4. The spacing between adjacent polymers can adjust to accommodate fused target proteins having a wide range of sizes. 5. Connections between the target protein and the TELSAM polymer need not be perfectly rigid because the resulting crystal lattice provides the remainder of the needed rigidity, forcing the target protein to choose among a small number of low energy orientations available to it.

Our ultimate goal is to develop a protein crystallization chaperone that consistently meets the following requirements: 1. Is easy to express in E. coli and purify with a yield greater than 5 mg of protein per liter of cell culture, even when fused to target proteins. 2. Enables fused targets to be soluble to at least 20 mg/mL. 3. Crystallizes within 30 days. 4. Forms crystals that are > 100 μm along their major axis and that diffract to better than 3 Å with an estimated mosaicity < 2°. 5. Results in target proteins being well-resolved in the crystallographic lattice following molecular replacement.

All published crystal structures of TELSAM alone or genetically fused to target proteins at the time of this writing feature direct inter-polymer contacts (**Figure 1A–F**)[5,6,7]. This observation led us and others [6] to hypothesize that strong inter-polymer contacts are essential to obtain well-diffracting crystals of TELSAM–target protein fusions. Previously-reported TELSAM–target fusions utilized 2TEL (which fuses 2 copies of the TEL-SAM domain in tandem and thus displays 3 copies of the target protein around the 6-fold TELSAM polymer axis) or 3TEL (which fuses 3 copies of the TEL-SAM domain in tandem and thus displays 2 copies of the target protein around the TELSAM polymer axis)[6, 7](**Figure 1G**). Both of these architectures allowed direct inter-polymer contacts while also accommodating the target protein in the crystal lattice (**Figure 1D-F**). We hypothesized that fusion of a target protein to 1TEL (which would display 6 copies of the target protein around the TELSAM polymer axis) would prevent TELSAM from making any inter-polymer contacts. Instead, all crystal contacts would have to be made by the fused target protein (**Figure 1H**). We further hypothesized that the inability to make direct inter-polymer contacts might preclude 1TEL–target fusions from forming well-diffracting crystals. We also hypothesized that fusion of a target protein to 3TEL (displaying 2 copies of the target protein around the TELSAM polymer axis) would allow at least some direct inter-polymer contacts and more readily form well-diffracting crystals (**Figure 1I**). We report the results of testing this hypothesis.

**Figure 1:**
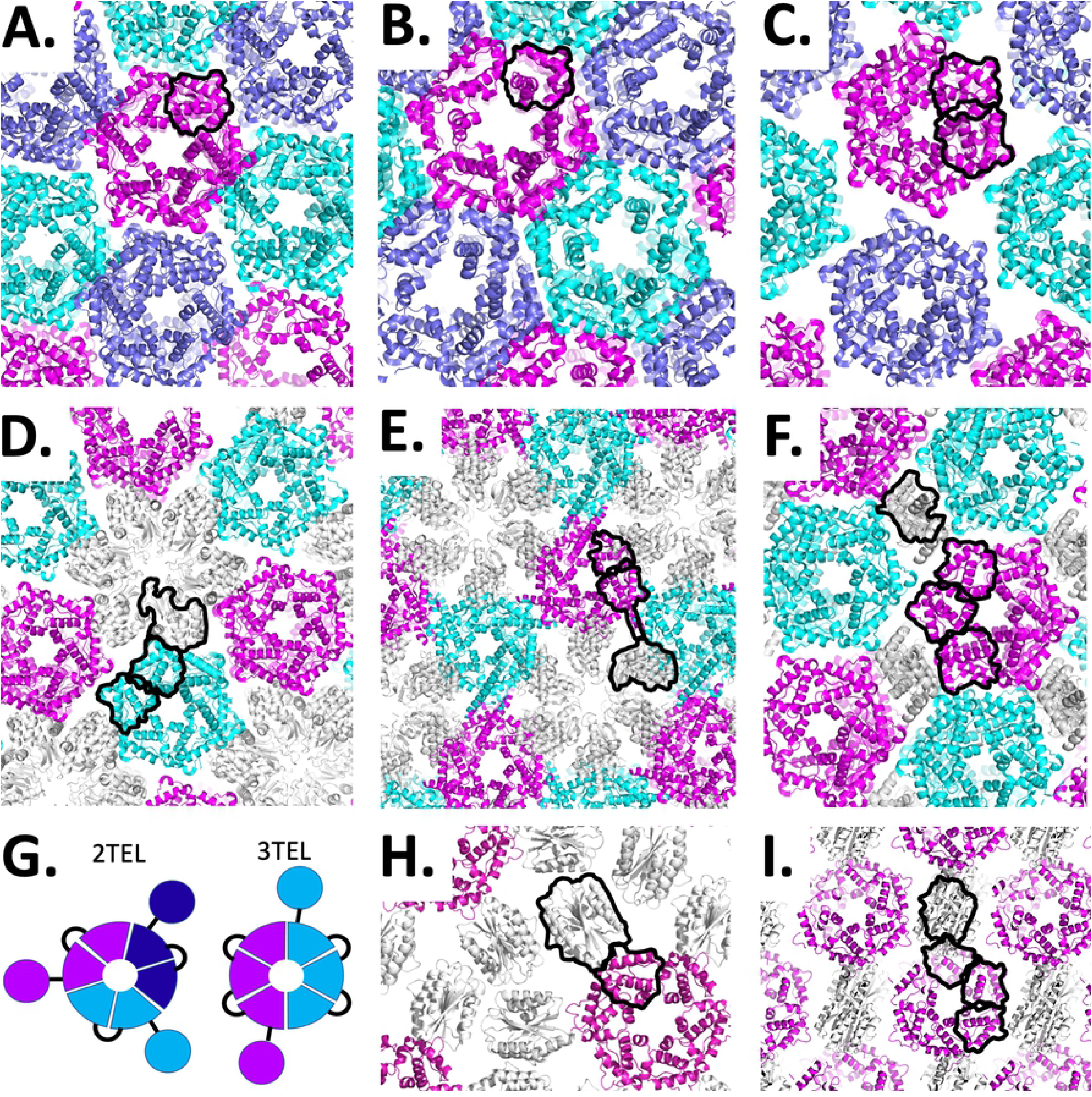
All currently-reported structures involving TELSAM polymers feature direct inter-polymer crystal contacts. TELSAM polymers are shown in cartoon representation and colored in magenta, cyan, and slate. Fused target proteins are colored gray. In each image, a single polypeptide within a single polymer has been indicated with black outlines around each of its constituent sub-domains. A. PDB ID 1JI7: 1TEL alone, B. PDB ID 1LKY: 1TEL E222R mutant, C. PDB ID 2QB1: 2TEL alone, D. PDB ID 2QB0: 2TEL-Lysozyme fusion, E. PDB ID 2QAR: 2TEL-helix-Lysozyme fusion, F. PDB ID 5L0P: 3TEL-ferric uptake regulator fusion, G. Schematic of 2TEL and 3TEL. Individual polypeptides are offset with unique colors. Circles denote target proteins, wedges denote TELSAM subunits, and black lines denote linkers. H. Potential crystal packing of a 1TEL-target protein fusion. I. Potential crystal packing of a 3TEL-target protein fusion.

## Materials and Methods

### Cloning the vWa alone

Residues 38 to 217 from human ANTXR cell adhesion molecule 2 (ANTXR2)(also known as capillary morphogenesis gene 2 (CMG2))(Uniprot: P58335) comprising the von Willebrand Domain (vWa) were reverse-translated, codon optimized (DNAworks), and synthesized as a gene fragment (Twist Biosciences). The 2 cysteines in this region of the gene were first mutated to alanine. The gene fragment was cloned into a custom pET42_SUMO vector using Gibson assembly [8], transformed into BL21(DE3) cells, and sequence-verified. pET42_SUMO was derived from pET42 by inserting a 10xHis–Yeast SMT3–XhoI fragment between its NdeI and AvrII sites, in place of the GST gene (Novagen).

### 10xHis-SUMO-CMG2 amino acid sequence (dots offset domains and linkers)

MGHHHHHHHHHHGS…LQDSEVNQEAKPEVKPEVKPETHINLKVSDGSSEIFFKIKKTTPLRRLMEAFAKRQGKEMDSLRFLYDGIRIQADQAPEDLDMEDNDIIEAHREQIGG…SARRAFDLYFVLDKSGSVANNWIEIYNFV QQLAERFVSPEMRLSFIVFSSQATIILPLTGDRGKISKGLEDLKRVSPVGETYIHEGLKLANEQIQKAGGLKTSSIIIALTDGKLDGLVPSYAEKEAKISRSLGASVYAVGVLDFEQAQLERIADSKEQVFPVKGGFQALKGIINSILAQS

### Production and crystallization of the vWa domain

A stab of frozen cell stock was used to inoculate 60 mL of LB media supplemented with 0.35 % glucose and 100 μg/mL kanamycin. This was shaken overnight at 30°C and 250 RPM. The next day, 10 mL of this overnight culture was used to inoculate 1 L of LB media supplemented with 0.05% glucose and 100 μg/mL kanamycin. This was also shaken at 37°C and 250 RPM. At an OD of 0.5, isopropyl β-D-1-thiogalactopyranoside (IPTG) was added to a final concentration of 0.1 mM to the culture. The culture was cooled to 18°C and shaken at 250 RPM for an additional 20 hours. The cells were collected by centrifugation, snap-frozen in liquid nitrogen, and stored at - 80°C.

All purification steps were completed on ice or in a 4°C refrigerator. 20 g of wet cell paste was resuspended in 100 mL of wash buffer (50 mM Tris, pH 7.3, 200 mM KCl, 50 mM imidazole, and 10 mM MgCl_2_), supplemented with 1 mM phenylmethylsulfonyl fluoride (PMSF) and 100 μM dithiothreitol (DTT). Lysozyme, deoxyribonuclease I, and ribonuclease were added to the resuspended cells to final concentrations of 20 μM, 800 nM, and 2 μM, respectively, and the lysis reaction was sonicated for 25 cycles of 12 seconds on at 59 seconds off at 60% power (Qsonica Q500) in a spinning ice bath. The lysate was clarified by centrifugation and applied to three 3 mL pre-equilibrated Ni-NTA columns, which were then washed with 5 CV of wash buffer. The protein was eluted using 35 CV of elution buffer (50 mM Tris, pH 7.3, 200 mM KCl, 400 mM imidazole, and 10mm MgCl_2_) and desalted using several PD-10 columns in parallel (Cytiva). Approximately 50 mg of pure protein was obtained per liter of culture. The protein was combined with 2 mg of purified SUMO protease [9] and DTT to a final concentration of 0.1 mM. The cleavage reaction was incubated overnight at 4°C. The following day, the cleavage reaction was applied to 2 mL of fresh Ni-NTA resin (to capture the SUMO protease, cleaved 10x-His-SUMO tags, and any uncleaved protein). After cleavage, the protein was diluted 8-fold with water and applied to a 4 mL Source 15Q anion exchange column (Cytiva). The vWa eluted in the flow through, while contaminants bound the resin and eluted in a KCl gradient. The vWa domain was further purified by size exclusion chromatography using a 100 mL Superdex 200 prep grade column.

Following SEC, the protein was buffer exchanged into 12.5 mM Tris, pH 7.3, 200 mM KCl. 1.2 μL of 20, 30, or 40 mg/mL vWa was combined with 1.2 μL of reservoir solution in a sitting drop format (SPT Labtech Mosquito). Commercially available crystallization screens: PEG Ion, Index, Salt-Rx, PEG-Rx (Hampton Research) and custom screens (PEG-custom, Bis-Tris Mg-Formate, PEG-KSCN, PEG-malonate) were employed. 82 days after setting the trays, a single rhomboid crystal (200 x 200 x 200 μm) appeared in 100 mM glycine, pH 9.5, 30% PEG 3350.

X-ray diffraction data was collected remotely at SSRL beamline 9-2. This crystal diffracted to around 2.1 Å resolution (I/σ > 2) but was highly mosaic, exhibiting spot splitting. The top indexing solution only included 9% of the reflections, complicating space group assignment. The data was processed using the Autoproc pipeline [10] and the Staraniso algorithm [11]. The phases were solved by molecular replacement using Phenix Phaser [12, 13] The structure then went through alternating stages of rebuilding in Coot [14] and refinement in Phenix Refine [15, 16]. TLS parameters were refined as well, using TLS groups calculated by Phenix. Refinement was assisted using statistics from the MolProbity server [17]. In spite of extensive refinement into P1, P2, P21212, and P212121 unit cells, acceptable R-factors could not be obtained.

### Cloning of 1TEL-flex-vWa

A gene fragment that placed residues 40 to 217 from human ANTXR cell adhesion molecule 2 (the vWa domain)[18] after residues 47 to 124 (the sterile alpha motif (SAM) domain) of human ETS variant transcription factor 6 (also known as Translocation ETS Leukemia (TEL))(Uniprot: P41212) was designed. TEL–SAM arginine 49 was mutated to alanine to alleviate a potential clash with the vWa domain. Other mutations in this gene relative to the human sequence were valine 112 to alanine and lysine 122 to alanine. A single alanine linker was placed between the TELSAM and the vWa domain, all cysteines were mutated to alanines, and vWa arginine 41 was mutated to alanine to alleviate a potential clash with TELSAM. The resulting amino acid sequence was reverse-translated, codon optimized (DNAworks) and synthesized by a commercial vendor (Integrated DNA Technologies). This gene fragment was cloned into the pET42-SUMO vector using Gibson assembly [8], transformed into BL21(DE3) cells, and sequence-verified (Eton Biosciences). Overlap PCR mutagenesis and Gibson assembly were then used to change the TELSAM alanine 112 to glutamate to make polymer formation triggerable by a reduction in pH, as previously described [5]. This construct was also transformed into BL21(DE3) cells and sequence-verified.

### 10xHis-SUMO-1TEL-flex-vWwa amino acid sequence

MGHHHHHHHHHHGS…LQDSEVNQEAKPEVKPEVKPETHINLKVSDGSSEIFFKIKKTTPLRRLMEAFAKRQGKEMDSLRFLYDGIRIQADQAPEDLDMEDNDIIEAHREQIGG…GSIALPAHLRLQPIYWSRDDVAQWLKWA ENEFSLRPIDSNTFEMNGKALLLLTKEDFRYRSPHSGDELYELLQHILAQ…A…RAAFDLYFVLDKSGSVANNW IEIYNFVQQLAERFVSPEMRLSFIVFSSQATIILPLTGDRGKISKGLEDLKRVSPVGETYIHEGLKLANEQIQKAG GLKTSSIIIALTDGKLDGLVPSYAEKEAKISRSLGASVYAVGVLDFEQAQLERIADSKEQVFPVKGGFQALKGIIN SILAQS

### Production and crystallization of 1TEL-flex-vWa

A stab of frozen cell stock was used to inoculate 40 mL of LB media, supplemented with 0.35% glucose and 100 μg/mL kanamycin. This was shaken overnight at 30°C and 250 rpm. The following day, 10 mL of the overnight culture was diluted into 1 L of LB media, supplemented with 0.05% glucose and 100 μg/mL kanamycin. This was again shaken at 37°C and 250 rpm. At an O.D. of 0.5, IPTG was added to a final concentration of 100 μM. The culture was cooled to 18°C and shaken at 250 RPM for an additional 20 hours. The cells were collected by centrifugation, snap-frozen in liquid nitrogen, and stored at -80°C.

All purification steps were completed on ice or in a 4°C refrigerator. 5 g of wet cell paste were resuspended in 25 mL of wash buffer (50 mM Tris, pH 8.8, 200 mM KCl, 50 mM imidazole, 10 mM MgCl_2_), supplemented with 1 mM PMSF, 100 μM DTT, 0.5 mg/mL lysozyme, and approximately 1 mg DNase. The cells were lysed by sonication for 25 cycles of 12 seconds on, 59 seconds off at 60% power (Qsonica Q500) in a spinning ice bath. The resulting lysate was clarified by centrifugation and applied to 2 mL of Ni-NTA resin, which was then washed with about 7 CV of wash buffer. The protein was then eluted with about 7 CV of elution buffer (50 mM Tris, pH 8.8, 200 mM KCl, 400 mM imidazole, 10 mM MgCl_2_) and desalted using several PD-10 desalting columns in parallel (Cytiva). Approximately 117 mg of protein was obtained per L of culture. The SUMO tag was removed by the addition of about 10 mg of SUMO protease [9] and DTT to 0.1 mM. The cleavage reaction was allowed to proceed overnight at 4°C. The SUMO tags and SUMO protease were removed by passing the protein solution over another 2 mL Ni-NTA column. The protein was then diluted 8-fold with water and applied to a 4 mL CaptoQ anion exchange column (Cytiva). The protein bound the column and was eluted in a KCl gradient. The protein was further purified by size exclusion chromatography using a 100 mL Superdex 200 prep grade column.

Following SEC, the protein was buffer exchanged into 12.5 mM Tris, pH 8.8, 200 mM KCl, and PMSF, phosphoramidon, and pepstatin A were added to final concentrations of 1 mM, 25 μM, and 1 μM, respectively. 1.2 μL of 20 mg/mL protein was combined with 1.2 μL of reservoir solution in a sitting drop format (SPT Labtech Mosquito). Commercially available crystallization screens (PEG Ion and Index [Hampton Research]) and custom screens (PEG-Tacsimate and PEG-Malonate) were employed. Several large crystals (100 x 100 x 500 μm) of 1TEL-flex-vWa appeared in 3 days in 100 mM Bis-Tris, pH 5.7, 3.0 M NaCl.

X-ray diffraction data was collected remotely at SSRL beamline 9-2. These crystals diffracted to around 2.8 Å resolution (I/σ > 2), were readily indexed in a primitive hexagonal unit cell with dimensions a = b = 104.8 Å, c = 57.6 Å, and exhibited low estimated mosaicity (0.4–0.6°). The data was processed using the Autoproc pipeline [10] and the staraniso algorithm [11]. The phases were solved by molecular replacement using Phenix Phaser [12, 13]. The structure then went through alternating stages of rebuilding in Coot [14] and refinement in Phenix Refine [15, 16]. TLS parameters were refined as well, using TLS groups found by the TLSMD server [19] Refinement was assisted using statistics from the MolProbity server [17].

### Cloning of 3TEL-flex-vWa

The 3TEL-flex-vWa gene was constructed by separately amplifying the 2TEL portion or the C-terminal 1TEL–vWa portion of a previously synthesized 2TEL-flex-vWA gene (Twist Bioscience). this template 2TEL-flex-vWa gene featured 2 TELSAM domains similar to that of the 1TEL-flex-vWa construct, fused using the same inter-TELSAM linker as PDB ID 2QAR [6], and had a vWa domain fused to the C-terminus of the second TEL–SAM domain using the same single alanine linker as for 1TEL-flex-vWa. The 2TEL and 1TEL-vWa PCR products were then fused by PCR and cloned into the pET42-SUMO vector using a modified Gibson assembly protocol [8]. The resulting plasmid was then transformed into BL21(DE3) E. coli cells and sequence verified (Eton Bioscience).

### 10xHis-SUMO-3TEL-flex-vWa amino acid sequence

MGHHHHHHHHHHGS…LQDSEVNQEAKPEVKPEVKPETHINLKVSDGSSEIFFKIKKTTPLRRLMEAFAKRQGKEMDSLRFLYDGIRIQADQAPEDLDMEDNDIIEAHREQIGG…GSIALPAHLRLQPIYWSRDDVAQWLKWA ENEFSLRPIDSNTFEMNGKALLLLTKEDFRYRSPHSGDELYELLQHILKQRPGGGGST…SIALPAHLRLQPIYWS RDDVAQWLKWAENEFSLRPIDSNTFEMNGKALLLLTKEDFRYRSPHSGDVLYELLQHILKQRPGGGGST…SI ALPAHLRLQPIYWSRDDVAQWLKWAENEFSLRPIDSNTFEMNGKALLLLTKEDFRYRSPHSGDVLYELLQHIL KQ…ARAAFDLYFVLDKSGSVANNWIEIYNFVQQLAERFVSPEMRLSFIVFSSQATIILPLTGDRGKISKGLEDLKRVSPVGETYIHEGLKLANEQIQKAGGLKTSSIIIALTDGKLDGLVPSYAEKEAKISRSLGASVYAVGVLDFEQAQLERIADSKEQVFPVKGGFQALKGIINSILAQS

### Production and crystallization of 3TEL-flex-vWa

A stab of frozen cell stock was used to inoculate 60 mL of LB media, supplemented with 0.35% glucose and 100 μg/mL kanamycin. This was shaken overnight at 30°C and 250 rpm. The following day, 10 mL of the overnight culture was diluted into 1 L of LB media, supplemented with 0.05% glucose and 100 μg/mL kanamycin. This was also shaken at 37°C and 250 rpm. At an O.D. of 0.5, IPTG was added to a final concentration of 0.1 mM. The cultures were next incubated for an additional 2 hours at 37°C with shaking at 250 rpm. The cells were then centrifuged, collected, frozen in liquid nitrogen, and stored at -80°C.

All purification steps were completed on ice or in a 4°C refrigerator. 20 g of wet cell paste were resuspended in 100 mL of wash buffer (50 mM Tris, pH 8.8, 200 mM KCl, 50 mM imidazole, 10 mM MgCl2) and supplemented with 1 mM PMSF, 100 μM DTT, 0.5 mg/mL lysozyme, and approximately 1 mg DNase. The cells were lysed by sonication for 25 cycles of 12 seconds on, 59 seconds off at 60% power (Qsonica Q500) in a spinning ice bath. The resulting lysate was clarified by centrifugation and applied to 2 mL of Ni-NTA resin which was next washed with 7 CV of wash buffer. The protein was then eluted with 7 CV of elution buffer (50 mM Tris, pH 8.8, 200 mM KCl, 400 mM imidazole, 10 mM MgCl2) and desalted using several PD-10 desalting columns in parallel (Cytiva). Approximately 63 mg of protein was obtained per L of culture. The SUMO tag was removed by the addition of 2 mg of SUMO protease [9] and DTT to 0.1 mM. The cleavage reaction was allowed to proceed overnight. The SUMO tags and SUMO protease were removed from the protein solution by passing it over another 2 mL of Ni-NTA column. The protein was diluted 8-fold with water to bring the KCl concentration to 25 mM and applied to a 4 mL Source 15Q anion exchange column. The 3TEL-flex-vWa did not bind the resin, while other impurities did and were thus removed. The protein further purified using a 100 mL Superdex 200 prep grade column.

The pure protein was concentrated to 20 mg/mL and screened against commercially available crystallization screens (Hampton Research: PEG Ion, Index, Salt-Rx, PEG-Rx) and in-house custom screens (PEG-custom, Bis-Tris Mg Formate) in a sitting drop format at room temperature. Rod-like crystals of 3TEL-flex-vWa also appeared within 7 days in many conditions, with the largest (50 x 50 x 200 μm) appearing in 0.1 M ammonium tartrate, pH 6.8, 12 % w/v PEG 3350.

X-ray diffraction data was collected remotely at SSRL beamline 9-2. These crystals diffracted to around 2.7 Å resolution but gave unusual diffraction patterns and exhibited translational pseudosymmetry. The data was processed using the Autoproc pipeline [10] and the Staraniso algorithm [11]. Molecular replacement was attempted using Phenix Phaser [12, 13].

### Cloning of 3TEL-rigid-DARPin

The 3TEL-rigid-DARPin (<u>D</u>esigned <u>A</u>nkyrin <u>R</u>epeat <u>P</u>rote<u>in</u>) construct was synthesized by a commercial vendor (Twist Bioscience), inserted into the pET42_SUMO vector, transformed into BL21(DE3) cells, and sequence-verified.

### 10x-His-SUMO-3TEL-rigid-DARPin amino acid sequence

MGHHHHHHHHHHGS…LQDSEVNQEAKPEVKPEVKPETHINLKVSDGSSEIFFKIKKTTPLRRLMEAFAKRQGKEMDSLRFLYDGIRIQADQAPEDLDMEDNDIIEAHREQIGG…GSIRLPAHLRLQPIYWSRDDVAQWLKWA ENEFSLRPIDSNTFEMNGKALLLLTKEDFRYRSPHSGDELYELLQHILKQR…PGGGGST…SIRLPAHLRLQPIY WSRDDVAQWLKWAENEFSLRPIDSNTFEMNGKALLLLTKEDFRYRSPHSGDVLYELLQHILKQR…PGGGGS T…SIRLPAHLRLQPIYWSRDDVAQWLKWAENEFSLRPIDSNTFEMNGKALLLLTKEDFRYRSPHSGDVLYELL QHILKQR…DLEAEAAAAE…DLGKKLLEAARAGQDDEVRILMANGADVNATDNDGYTPLHLAASNGHLEIVEVLLKNGADVNASDLTGITPLHAAAATGHLEIVEVLLKHGADVNAYDNDGHTPLHLAAKYGHLEIVEVLLKHGADVNAQDKFGKTAFDISIDNGNEDLAEILQKLN

### Production and crystallization of 3TEL-rigid-DARPin

A stab of frozen cell stock was used to inoculate 40 mL of LB media, supplemented with 0.35% glucose and 100 μg/mL kanamycin. This was shaken overnight at 30°C and 250 rpm. The following day, 10 mL of the overnight culture was diluted into 1 L of LB media, supplemented with 0.05% glucose and 100 μg/mL kanamycin. This was again shaken at 37°C and 250 rpm. At an O.D. of 0.5, IPTG was added to a final concentration of 100 μM. The culture was cooled to 18°C and shaken at 250 RPM for an additional 24 hours. The cells were collected by centrifugation, snap-frozen in liquid nitrogen, and stored at -80°C

All purification steps were completed on ice or in a 4°C refrigerator. 25 g of wet cell paste was resuspended in 125 mL of wash buffer (50 mM Tris, pH 8.8, 200 mM KCl, 50 mM imidazole), supplemented with 1 mM PMSF, 100 μM DTT, and approximately 1 mg DNase. The cell suspension was homogenized at 18,000 psi for 2 passes (NanoDeBEE, BEE International). The resulting lysate was clarified by centrifugation at 40000 x g and applied to 2 mL of HisPure Ni-NTA resin, which was then washed with 7 CV of wash buffer. The protein was then eluted with 7 CV of elution buffer (50 mM Tris, pH 8.8, 200 mM KCl, 400 mM imidazole, 10 mM MgCl2) and desalted using several PD-10 desalting columns in parallel (Cytiva). Approximately 57 mg of protein was obtained per L of culture. The SUMO tag was removed by the addition of 30 mg of SUMO protease [9] and DTT to 0.1 mM. The cleavage reaction was allowed to proceed overnight at 4°C. The SUMO tags and SUMO protease were removed from the protein solution by passing it over another 2 mL Ni-NTA column. The protein was diluted 8-fold with water and applied to a 4 mL Source 15Q anion exchange column (Cytiva). The protein bound the column and was eluted in a KCl gradient. The protein was further purified by size exclusion chromatography using a 100 mL Superdex 200 prep grade column.

Following SEC, the protein was buffer exchanged into 12.5 mM Tris, pH 8.8, 200 mM KCl, and PMSF, phosphoramidon, and pepstatin A were added to final concentrations of 1 mM, 25 μM, and 1 μM, respectively. 1.2 μL of 15 mg/mL protein was combined with 1.2 μL of reservoir solution in a sitting drop format (SPT Labtech Mosquito). Commercially available crystallization screens (PEG Ion, SaltRX, PEG-Rx, and Index [Hampton Research]) and custom screens (PEG-custom and Bis-Tris Mg Formate) were employed. Thin plate crystals (10 x 100 x 100 μm) appeared in 3 days in many conditions, with the largest (100 x 100 x 10 μm) in 0.2 M L-Proline, 0.1 M HEPES, pH 7.4, 10 % w/v PEG 3350 and in 0.05 M MgCl_2_, 0.1 M HEPES, pH 7.3, 30 % v/v PEG 550 MME.

X-ray diffraction data was collected remotely at SSRL beamline 9-2. A single thin plate crystal diffracted to 3.2 Å resolution (I/σ of 2.0)(SSRL beamline 9-2) and was indexed in a primitive monoclinic unit cell with dimensions a = 45.9 Å, b = 165.6 Å, c = 63.6 Å, α = 90°, β = 90.162°, γ = 90°. The top indexing solution only included 12% of the reflections. The data was processed using the Autoproc pipeline [10] and the Staraniso algorithm [11]. The phases were solved by molecular replacement using Phenix Phaser [12, 13] The structure then went through alternating stages of rebuilding in Coot [14] and refinement in Phenix Refine [15, 16]. Initially a P212121 space group was used with one 3TEL-rigid-DARPin molecule in the asymmetric unit, but a P1211 space group with 2 molecules in the asymmetric unit gave superior refinement statistics. TLS parameters were refined as well, using TLS groups found by the TLSMD server [19]. Refinement was assisted using statistics from the MolProbity server [17].

## Results

TELSAM greatly increases the crystallization rate of the human ANTXR cell adhesion molecule 2 vWa domain. The vWa domain was chosen because it has been successfully crystallized previously, has a known structure, and is known to be only moderately soluble. These properties make the vWa domain an excellent representative target protein to evaluate potential crystallization chaperones. We sought to quantitatively evaluate whether fusion to TELSAM could enhance the propensity or rate of vWa crystallization relative to the vWa alone. We produced soluble vWa domain and purified it using nickel-NTA, anion exchange chromatography, and size exclusion chromatography, achieving excellent purity. We then executed crystallization trials at a range of protein concentrations and using a range of commercially available and custom-made crystallization screens. 82 days after setting the trays, a large rhomboid crystal appeared, diffracting to around 2.1 Å resolution. This crystal was highly mosaic, exhibited spot splitting, and the top indexing solution only included 9% of the reflections, complicating space group assignment. Molecular replacement was carried out using a published structure of the vWa domain (PDB ID: 1SHU)[18] and a series of space group possibilities. In spite of extensive refinement into P1, P2, P21212, and P212121 unit cells, acceptable R-factors could not be obtained. The vWa crystal was not obtained until anion exchange chromatography was added to the purification protocol.

We next modeled the shortest flexible genetic fusion of the human ANTXR cell adhesion molecule 2 vWa domain structure [18] to the C-terminus of a single TELSAM monomer (1TEL)(ref) using PyMOL (Schrödinger) and Foldit [20]. We determined that the vWa domain could be flexibly fused to TELSAM with a linker consisting of a single alanine residue (**Figure 2A**). This construct included the critical TELSAM V112E mutation that makes TELSAM polymerization pH-dependent [5]. Two constructs were made. One fused the vWa domain to a single TELSAM monomer (1TEL), while the other fused the vWa domain to the last of three linked, tandem TELSAM monomers (3TEL)(**Figure 2B**). The 3TEL construct was designed by inserting the flexible linker of PDB ID 2QAR [6] between the C-terminus of one TELSAM monomer and the N-terminus of another and then repeating the process to obtain 3 TELSAM monomers in tandem in a single polypeptide (**Figure 2C**)[7]. As previously described, this 3TEL construct contained the V112E mutation only in the N-terminal-most TELSAM domain [7].

**Figure 2:**
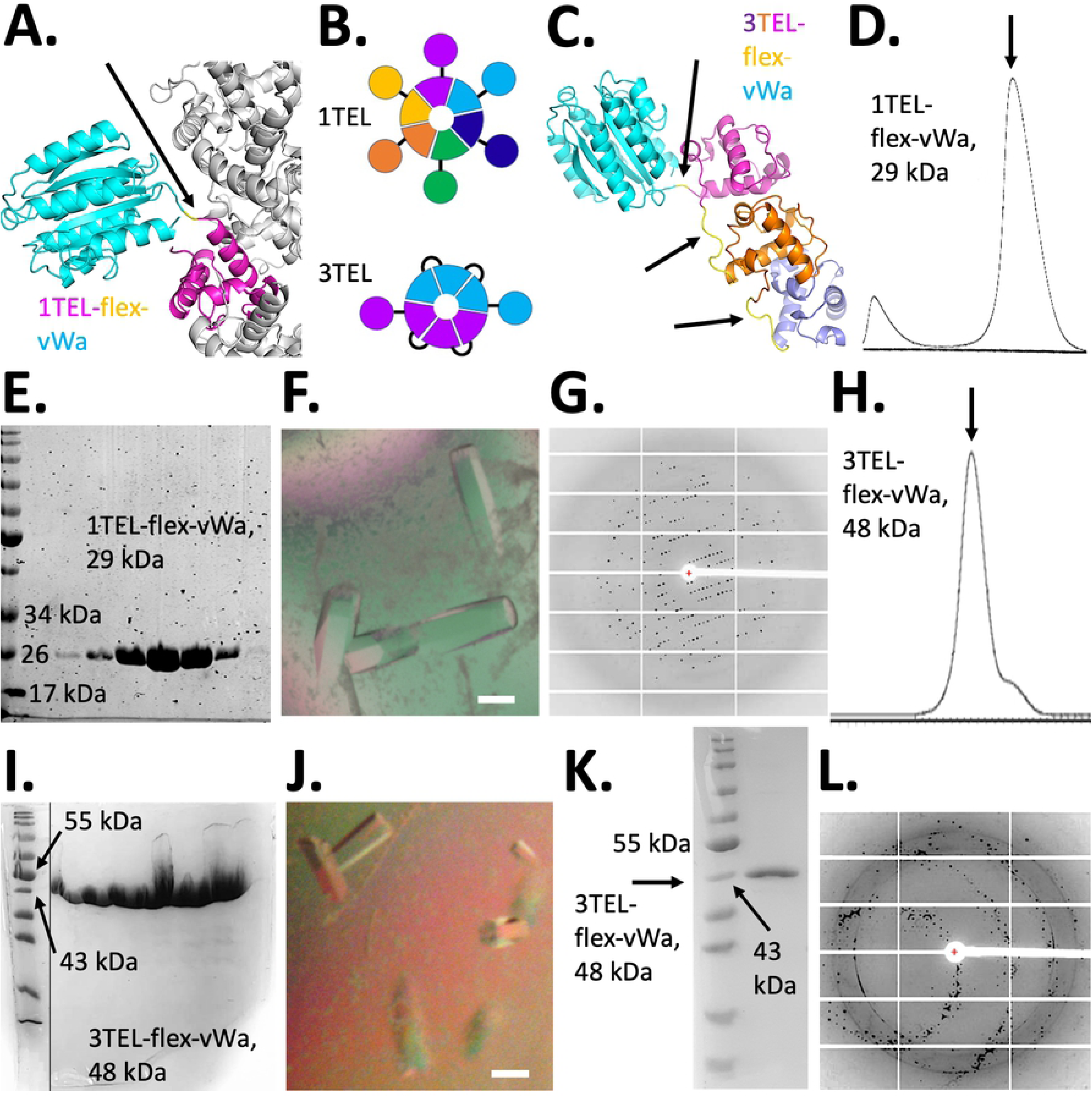
TELSAM greatly enhances the crystallization rate of genetically fused vWa domain. A. Design model of 1TEL-flex-vWa in cartoon representation, with TELSAM in magenta and vWa in cyan. Other subunits of the TELSAM polymer are shown in white. The alanine linker is indicated with an arrow. B. Schematic of 1TEL and 3TEL. Individual polypeptides are offset with unique colors. Circles denote target proteins, wedges denote TELSAM subunits, and black lines denote linkers. C. Detail of 3TEL-flex-vWa tandem fusion, with successive 1TEL domains shown in purple, orange, and magenta and the vWa domain in cyan. Linkers are indicated with arrows. D. SEC trace of 1TEL-flex-wWa. E. PAGE gel of purified 1TEL-flex-wWa. F. Crystals of 1TEL-flex-vWa, scale bar is 100 μm. G. Representative diffraction pattern from a crystal of 1TEL-flex-vWa. H. SEC trace of 3TEL-flex-wWa. I. PAGE gel of purified 3TEL-flex-wWa. The vertical black line indicates where less important lanes of the gel have been cropped out. J. Crystals of 3TEL-flex-vWa, scale bar is 100 μm. K. PAGE gel of washed 3TEL-flex-vWa crystals. L. Representative diffraction pattern from a crystal of 3TEL-flex-vWa.

The resulting proteins were produced, purified, and crystallized with in a manner identical to the solitary vWa domain above (**Figure 2D,E,H,I**), except that a cocktail of protease inhibitors were added to the pure protein immediately before setting crystallization drops (to prevent cleavage of the TELSAM–TELSAM or TELSAM–vWa linkers by trace proteases). Large crystals of 1TEL-flex-vWa appeared in 3 days (**Figure 2F**), diffracting to around 2.8 Å resolution (**Figure 2G, Table 1**). As with the vWa alone, crystals of 1TEL-flex-vWa were not obtained until anion exchange chromatography was added to the purification protocol. Notably, 1TEL-flex-vWa could be successfully produced and crystallized by at least 3 independent teams of students in our research group.

**Table 1.** Crystallographic Data Collection and Refinement Statistics

| Construct | 1TEL-flex-vWa | 3TEL-rigid-DARPin |
| --- | --- | --- |
| PDB ID | 7N1O | 7N2B |
| **Data Collection** |  |  |
| X-ray source | SSRL 9-2 | SSRL 9-2 |
| Wavelength | 0.979460 | 0.979460 |
| Detector Type | Pilatus 6M PAD | Pilatus 6M PAD |
| Detector Distance (mm) | 350 | 350 |
| Resolution range | 38.16-2.77 (2.869-2.77) | 40.16 - 3.221 (3.336 - 3.221) |
| Space group | P 65 | P 1 21 1 |
| Cell a, b, c (Å) | 103.396, 103.396, 56.55 | 45.962, 63.625, 166.005 |
| Cell $\alpha$, $\beta$, $\gamma$ (°) | 90, 90, 120 | 90, 90.162, 90 |
| Total reflections | 47554 (5106) | 46124 (4911) |
| Unique reflections | 8679 (878) | 13842 (1491) |
| Multiplicity | 5.5 (5.8) | 3.3 (3.3) |
| Completeness (%) | 97.10 (97.99) | 87.26 (95.36) |
| I/σI | 18.35 (2.88) | 10.98 (2.46) |
| Wilson B-factor | 71.05 | 84.40 |
| R-merge | 0.05193 (0.6253) | 0.07087 (0.5057) |
| R-meas | 0.05753 (0.6876) | 0.08449 (0.6052) |
| R-pim | 0.024 (0.2779) | 0.04539 (0.3275) |
| CC1/2 | 0.999 (0.899) | 0.999 (0.906) |
| CC* | 1 (0.973) | 1 (0.975) |
| **Refinement** |  |  |
| Reflections used in refinement | 8667 (878) | 13776 (1480) |
| Reflections used for R-free | 432 (44) | 681 (69) |
| R-work | 0.2030 (0.3449) | 0.2262 (0.3233) |
| R-free | 0.2285 (0.3904) | 0.2425 (0.3735) |
| CC (work) | 0.965 (0.802) | 0.956 (0.849) |
| CC (free) | 0.942 (0.519) | 0.969 (0.720) |
| Number of non-H atoms | 1932 | 6067 |
| Macromolecule atoms | 1921 | 6067 |
| Ligands | 3 | 0 |
| Water | 8 | 0 |
| Protein residues | 257 | 806 |
| Bond length R.M.S. Deviations | 0.009 | 0.003 |
| Bond angle R.M.S. Deviations | 0.86 | 0.56 |
| Ramachandran Stats (%) |  |  |
| Favored (%) | 98.04 | 98.36 |
| Allowed (%) | 1.96 | 1.64 |
| Outliers (%) | 0.00 | 0.00 |
| Rotamer outliers (%) | 0.00 | 0.00 |
| Clashscore | 8.23 | 4.73 |
| B-factors (Å<sup>2</sup>) |  |  |
| Average B-factor | 75.23 | 92.51 |
| Macromolecules | 75.26 | 92.51 |
| Ligands | 92.60 | N/A |
| Water | 61.89 | N/A |
| Number of TLS groups | 4 | 6 |

Statistics for the highest-resolution shell are shown in parentheses.

Rod-like crystals of 3TEL-flex-vWa also appeared within 7 days in various conditions (**Figure 2J**) and diffracted to around 2.7 Å resolution. These crystals contained highly pure protein of the correct molecular weight (**Figure 2K**) but gave unusual, smeared diffraction patterns at certain diffraction angles (**Figure 2L**). These datasets also exhibited translational pseudosymmetry and significant anisotropy in the diffraction limits (a = 2.154 Å, b= 2.154 Å, c = 3.792 Å, assuming a P6 space group). Thus far we have been unable to produce a molecular replacement solution with acceptable refinement statistics. Studies with 3TEL-flex-vWa are ongoing. The crystallization time and propensity of each of the vWa-containing constructs is summarized in **Table 2**.

**Table 2.** Crystallization Time, Propensity, and Diffraction quality of vWa Constructs

| Construct | Days to crystal appearance | Number of initial screening conditions producing crystals | Percent of reflections indexed |
| --- | --- | --- | --- |
| vWa alone | 82 | 1 | 9% |
| 1TEL-flex-vWa | 3 | 1 | 67% |
| 3TEL-flex-vWa | 7 | 19 | 50% |

Molecular replacement with the 1TEL-flex-vWa datasets was carried out by separately placing the structures of 1TEL (PDB ID: 2qb1)[6] and the vWa domain (PDB ID: 1shu)[18]. We determined the space group to be P65 (as expected due to the 6-fold symmetric, left-handed helical nature of the 1TEL polymer), with 1 molecule of 1TEL-flex-vWa per asymmetric unit. In the 1TEL-flex-vWa crystallographic lattice, the vWa domains make a head to head interaction, leaving considerable aqueous space (59.6% solvent content). Two molecules of the vWa domain separate adjacent TELSAM polymers and there are no direct inter-TELSAM polymer contacts, a feature never before seen in crystal structures of TELSAM-target protein fusions (**Figure 3A**). In spite of the lack of direct inter-polymer contacts and the high degree of aqueous space, crystals of 1TEL-flex-vWa exhibit minimal anisotropy, with the anisotropic diffraction limits of the unit cell axes being a = b = 2.60 Å, c = 2.55 Å [11]. Of note, we observed positive difference density extending away from the Mg^2+^ ion bound in the MIDAS site of the vWa domain. Based on the molecules present during protein purification and crystallization, we modelled a chloride ion into this density, coordinating the Mg^2+^ ion. The vWa from the crystal structure of 1TEL-flex-vWa differs from previously-published structures of the vWa only in the conformations of some of its surface loops (**Figure 3B**).

**Figure 3:**
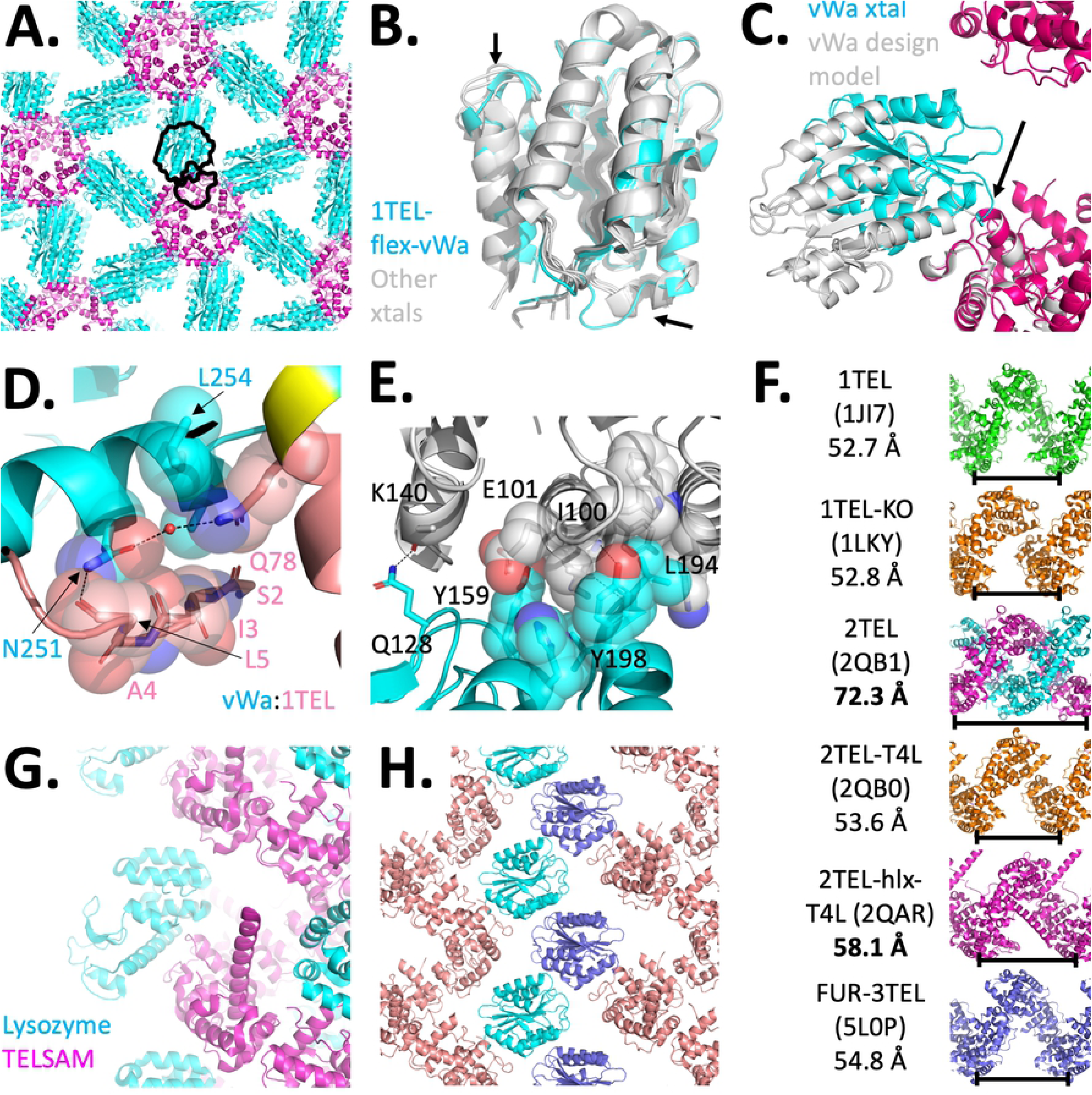
Detail of 1TEL-flex-vWa crystal structure and lattice. A. Crystal lattice of 1TEL-flex-vWa, in cartoon representation with TELSAM in magenta and the vWa in cyan. A black outline denotes each sub-domain of a single polypeptide within a single polymer. B. Superposition of the vWa domain from 1TEL-flex-vWa (magenta) with previously published vWa structures (gray, PDB IDs: 1SHU, 1SHT, 1T6B, 1TZN). C. Comparison of the design model (white) and crystal structure (magenta and cyan) of 1TEL-flex-vWa (the region of the linker that becomes α-helical is indicated with an arrow). Other copies of the vWa domain have been omitted for clarity. D. Detail of the *cis* interface between 1TEL (salmon) and vWa (cyan). Hydrogen bonds are shown as black dashes. The single alanine linker is shown in yellow. E. Detail of the *trans* interface between two vWa units, colored cyan and slate. F. Comparison of the helical rise from previously published TELSAM crystal structures. The relative rise of a single turn of each helix is denoted with a black bar. Fused target proteins have been omitted. G. Detail of the lysozyme (cyan) intercalation into the TELSAM polymer helix (magenta) of PDB ID: 2qar. H. Detail of the vWa domains (cyan and purple) zippering up and of their intercalation into the TELSAM polymer helix (salmon) of 1TEL-flex-vWa.

In the 1TEL-flex-vWa structure, the vWa unit adopts a position different from that of the design model, packing against the TELSAM polymer. The single amino acid alanine linker becomes an extension of the 1TEL C-terminal α-helix (**Figure 3C**). The C-terminal α-helix of the vWa domain packs against the N-terminus of 1TEL, burying 952 Å^2^ of solvent-accessible surface area (total of both sides of the interface) but making minimal interactions. Specifically, the vWa Leu 254 sidechain makes van der Waals interactions to the 1TEL Gln 78 side chain. The vWa Asn 251 sidechain amide carbonyl makes a direct hydrogen bond to the 1TEL Leu 5 main chain carbonyl and a water-mediated hydrogen bond to the 1TEL Gln 78 side chain amide nitrogen. The vWa Asn 251 side chain makes additional van der Waals contacts to the mainchain of 1TEL Ile 3 and the main chain and side chain of Ala 4. (**Figure 3D**). This demonstrates that flexibly-fused target proteins can find low-energy rigid conformations by packing against the TELSAM polymer when presented with the fast interaction on-rate conferred by covalent attachment.

The inter-vWa interface buries 982 Å^2^ of solvent-accessible surface area and is largely hydrophobic, with minimal polar interactions. Leu 194 from a vWa molecule packs into a hydrophobic pocket formed by the Trp 99, Ile 100, Tyr 103, and Lys 150 of a second vWa molecule. Additionally, Gln 128 and Tyr 198 from the first vWa molecule respectively make hydrogen bonds to the backbone carbonyl oxygen of Lys 240 and the backbone amide nitrogen of Ile 100 from the second vWa molecule. Finally, Tyr 159 from the first vWa molecule makes an anion-π interaction with the side chain carboxylate of Glu 101 and a cation-π interaction with the side chain carbonyl oxygen of Gln 104, both from the second vWa molecule. Conversely, Asn 104 and the Ile 100 from this second vWa molecule pack into a largely hydrophobic pocket formed by the His 161, Leu 194, Val 195, and Tyr 198 of the first vWa molecule. Asn 104 additionally makes a hydrogen bond to the backbone amide nitrogen of Leu 194 (**Figure 3E**).

This inter-vWa contact has not been observed in any previously-reported structure of the vWa domain [18,21,22]. In view of the fact that a flexibly-fused target protein can find a rigid conformation relative to the TELSAM polymer by packing against the polymer, the rigid hydrophobic connection between vWa domains suggest that while direct inter-polymer contacts are dispensable in forming well-diffracting crystals, a rigid transform between adjacent TELSAM polymers is still required.

Among previously-reported TELSAM structures, the TELSAM polymer helix has an average helical rise of 53.5 ± 0.7 Å (**Figure 3F**)[5,6,7]. Notable exceptions to this include the structure of 2TEL alone, which forms a double helix and so has a greatly expanded helical rise of 72.3 Å to accommodate the second intercalated helix [6]. Another exception is the structure of 2TEL fused to T4 lysozyme via a long α-helix, which has a slightly expanded helical rise of 58.1 Å, apparently due to partial intercalation of the lysozyme unit into the polymer helix (**Figure 3G**)[6]. The new structure of 1TEL-flex-vWa likewise has a slightly expanded helical rise of 56.9 Å, apparently due to the helical rise necessary to make optimal vWa-vWa contacts (**Figure 3H**). There is a possibility that the vWa domain perturbs the 1TEL helical rise because it intercalates into the 1TEL polymer helix, but this is unlikely because the vWa domains make no contacts to the next turn of the 1TEL polymer (the nearest resolved atoms of the vWa are at least 7.9 Å from the nearest atoms of the next turn of its own 1TEL polymer)(**Figure 3H**). A more likely explanation is that this degree of helical rise has been dictated by the spacing required to achieve the observed inter-vWa crystal contacts. The fact that the 1TEL helical rise is perturbed suggests that TELSAM is able to adjust its helical rise to accommodate the crystal packing interactions of fused target proteins. The 1TEL-flex-vWa structure also provides further evidence that the flexibility in the TELSAM helical rise is not necessarily a detriment to the growth of ordered crystals.

In related work, we modeled a rigid α-helical fusion between the C-terminal α-helix of 3TEL and the N-terminal α-helix of a DARPin (PDB ID 4J7W)[23] using PyMOL (Schrödinger). We chose a DARPin orientation that would allow it to non-covalently bind a second target protein using the DARPin’s canonical binding surface while minimizing clashes between that second target protein and the TELSAM polymer (**Figure 4A**). The 3TEL construct was designed as described above. The resulting protein was produced and crystallized in a manner identical to the 1TEL-flex-vWa construct above (**Figure 4B-C**), except that Mg^2+^ was omitted. Thin plate crystals appeared in 3 days in various conditions, diffracting to around 3.2 Å resolution (**Figure 4D-E**).

**Figure 4:**
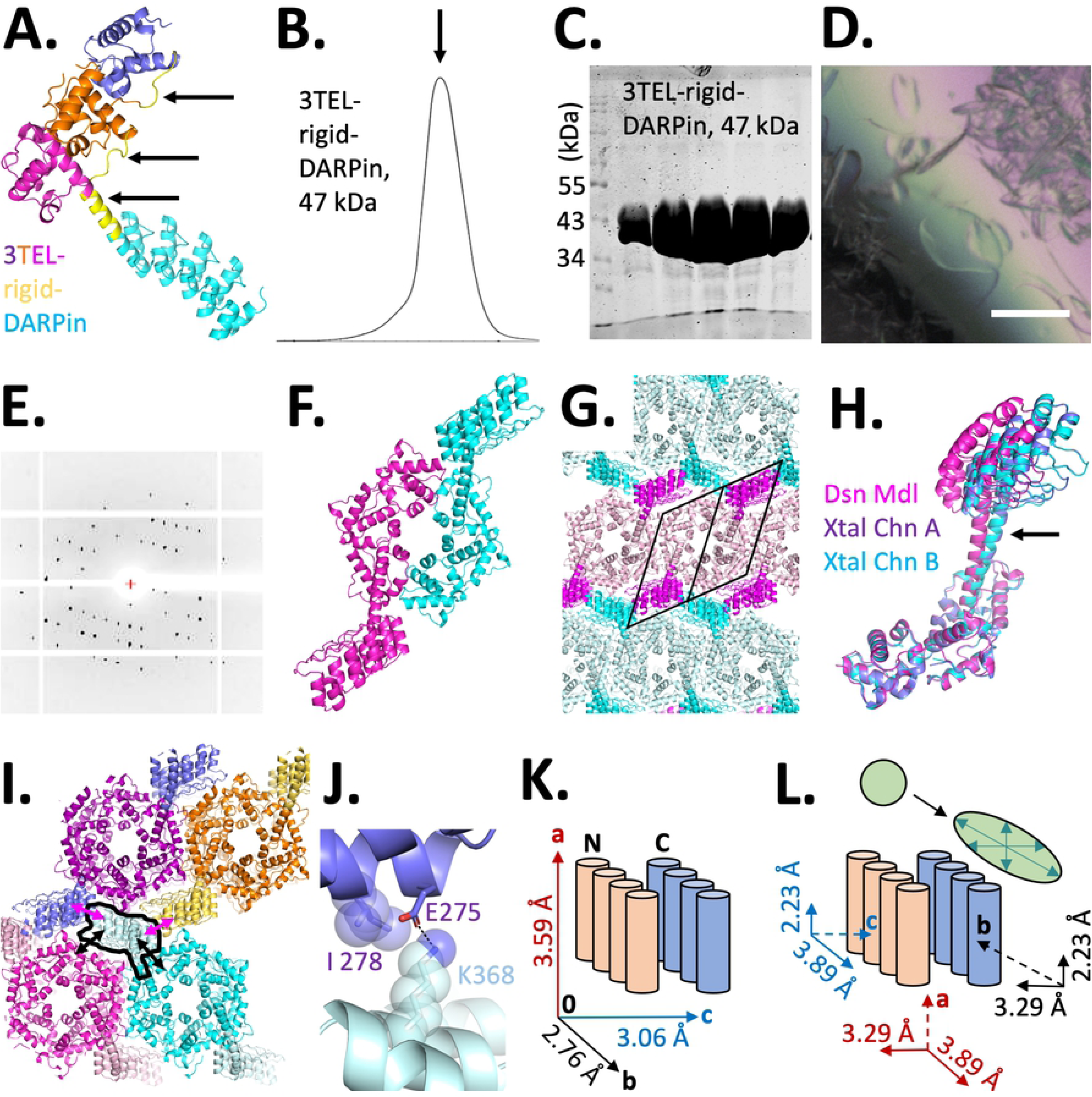
Production and detail of 3TEL-rigid-DARPin crystal structure. A. Design model of 3TEL-rigid-DARPin, with successive 1TEL domains shown in purple, orange, and magenta and the DARPin in cyan. Linkers are indicated with arrows. B. SEC trace of 3TEL-rigid-DARPin. C. Post-SEC PAGE gel of 3TEL-rigid-DARPin. D. Representative crystals of 3TEL-rigid-DARPin. Scale bar is 100 μm. E. Representative diffraction pattern of 3TEL-rigid-DARPin. F. Asymmetric unit, with 2 copies of 3TEL-rigid-DARPin. G. Crystal lattice of 3TEL-rigid-DARPin. A diamond denotes a single polymer while a parallelogram denotes a single polypeptide subunit. H. Comparison of design model (magenta) to crystal structure (cyan and slate) of 3TEL-rigid-DARPin, with an arrow to indicate the shift of the DARPin and connecting α-helix. I. Schematic of crystal contacts made by a single DARPin molecule, highlighted with a black line. Contacts to its own 3TEL polymer layer are indicated with black arrows, while contacts to DAPRins from an adjacent polymer layer are indicated with pink arrows. J. Interface between a DARPin (light blue) and another DARPin (slate) from an apposed 3TEL polymer layer, with selected amino acid side chains shown as sticks and a salt bridge shown as a black dash. This is the interface indicated with pink arrows in I. K. Schematic detailing the anisotropic diffraction limits of 3TEL-rigid-DARPin crystals along each unit cell axis. The orientation of two adjacent 3TEL polymer layers is denoted with colored cylinders. L. Schematic detailing how the anisotropic diffraction limits are decomposed into atomic displacements along each unit cell axis (solid line) when viewed along the orthogonal unit cell axis (dashed line). The observed diffraction limit is the average of the component atomic displacements. Also shown is an atom (green sphere) and a depiction of the positional uncertainty/atomic displacement of this atom (green oval) in the same unit cell frame of reference.

Molecular replacement into the 3TEL-rigid-DAPRin dataset was carried out by separately placing models of 3TEL and the DARPin (PDB ID: 4j7w)[23]. We determined the space group to be P1211, consistent with 2-fold symmetric, left-handed helical nature of the 3TEL polymer, with 2 molecules of 3TEL-rigid-DAPRin per asymmetric unit (**Figure 4F**). In the 3TEL-rigid-DARPin crystallographic lattice, 3TEL polymers pack side-by-side in lateral layers. All of the 3TEL polymers in a given layer are oriented in the same N-to-C direction but are oriented oppositely from the 3TEL polymers in the preceding or following layers. This is the first reported instance of a crystal structure that features TELSAM polymers not all aligned in the same N→C direction. Successive 3TEL layers are separated by a layer of DARPins that alternate emanating from one or the other of the layers (**Figure 4G**). The 3TEL-rigid-DARPin crystal structure differs from the design model only in that the DARPin translates as much as 12.5 Å in the crystal structure relative to the design model (**Figure 4H**). This confirms that an α-helical connection retains some residual flexibility, a feature observed in other studies utilizing rigid α-helical fusions [24,25,26]. This residual flexibility likely allowed the DARPin to access superior crystal packing arrangements.

3TEL-rigid-DARPin formed diffracting crystals with extremely minimal crystal contacts between adjacent layers of 3TEL-rigid-DARPin polymers. While a given DARPin makes a fair number of van der Waals and salt bridge contacts to its own 3TEL polymer (burying 462 Å of solvent-accessible surface area) and to an adjacent 3TEL polymer from the same polymer layer (burying 862 Å of solvent-accessible surface area), it makes no contacts to the 3TEL polymers of adjacent layers. All inter-layer contacts occur solely between DARPins from adjacent polymer layers (**Figure 4I**). As they are the only connections between adjacent 3TEL polymer layers, we were struck by how minimal the inter-DAPRin contacts were. Inter-DARPin contacts bury only 293 Å^2^ of solvent-accessible surface area and involve a single salt bridge between Lys 368 on the canonical binding surface of one DARPin and Glu 275 on the backside of a second DARPin. Ile 278 from the backside of the second DARPin also makes minimal van der Waals interactions to the Lys 368 (**Figure 4J**). Since this is a DARPin-DARPin contact, each DARPin makes 2 of these to other DARPins from the adjacent polymer layer. While the crystal packing of the vWa units slightly expanded the helical rise of the associated 1TEL polymer (**Figure 3H**), the DARPin crystal packing had a profound impact on the rise of the 3TEL polymer, reducing it to 45.9 Å, the minimum helical rise reported to date for a TELSAM polymer not involved in a double helix (**Figure 3F**). The DARPin units do not intercalate into the 3TEL polymer, ruling out target protein intercalation as a cause of the reduced degree of 3TEL helical rise, and confirming that inter-target crystal packing can dictate the rise of the TELSAM helical polymer.

The weak inter-DARPin crystal contacts correlate with the observed diffraction limits and scale factor of the thin axis of the crystal. Unlike the minimally anisotropic diffraction of 1TEL-flex-vWa crystals, 3TEL-rigid-DARPin crystals exhibited significant anisotropy as estimated using the Staraniso software [11]. The anisotropic diffraction limits of the 3TEL-rigid-DARPin unit cell were a = 3.59 Å, b = 2.76 Å, c = 3.06 Å (**Figure 4K,L**). When correlated with the unit cell axes, these anisotropic diffraction limits suggest that the largest displacement error is in the horizontal register of the helical 3TEL polymers (perpendicular to the helical axis of the polymers), either within each 3TEL layer or between successive layers. As the 3TEL polymers pack tightly against their neighbors within each polymer layer, we propose that the displacement error most likely lies in the horizontal register between successive 3TEL polymer layers and that successive polymer layers are able to shift horizontally relative to each other in the crystal (moving parallel to the plane of each polymer layer but perpendicular to the long axis of the individual polymers). This lateral shifting of polymer layers relative to each other may be a consequence of the extremely minimal inter-layer (inter-DARPin) crystal contacts (**Figure 4I,J**). Analysis of the crystal position during diffraction image collection reveals that the 3TEL polymer layers lie parallel to the long axes of the thin plate crystal. The thin axis of the crystal lies perpendicular to the planes of the 3TEL polymer layers and suggests that 3TEL-rigid-DARPin crystals experience a significant growth defect in this dimension. This growth defect may be due to the fact that successive 3TEL polymer layers contact the previous layer using only 2 glancing crystal contacts per DARPin. Concomitantly, diffraction images taken at angles close to parallel with the short c-axis of the crystal had a higher scale factor during data processing. That the c-axis of the unit cell lies parallel to the thin/short axis of the crystal could alternatively explain the lower resolution diffraction limits and higher scale factors of reflections collected from this angle.

## Discussion

In comparing the crystallization rate of the vWa domain from human CMG2, we have provided definitive evidence that a TELSAM-target protein fusion can crystallize more rapidly than the same target protein alone. We provide evidence to support the idea that fusion to TELSAM polymers confers immense avidity to fused target proteins to stabilize exquisitely weak crystal contacts made by the target proteins. We hypothesize that this is the principal manner in which TELSAM increases the crystallization propensity of fused target proteins. Based on the extremely weak individual inter-layer crystal contacts made in the 3TEL-rigid-DARPin structure, theoretically any sufficiently homogenous target protein should be readily crystallized through fusion to TELSAM. This observation strongly calls for continued investigation into the principles and requirements for reliably obtaining well-ordered crystals of TELSAM–target protein fusions.

We have shown that TELSAM-target protein fusions can form well-ordered, diffracting crystals when either 6 or 2 copies of the target protein are presented per turn of the TELSAM helical polymer. Future studies are underway to determine how generally useful fusion to TELSAM is for generating well-ordered crystals of a greater variety of target proteins, the optimal number of target proteins per turn of the TELSAM helical polymer, and whether this optimal number is dependent on the specific target protein. We hypothesize that 1TEL (with 6 copies of the target protein per helical turn of the polymer) is more likely to be generally successful because it has the potential to form triangular connections through the resulting crystal lattices (**Figure 1H, 3A**). These triangular connections are theoretically more resistant to distortion than the connections possible with 2TEL (hexagonal, 3 copies of the target protein per helical turn) and 3TEL (parallelogram-like, 2 copies of the target protein per helical turn), respectively (**Figure 1G,H,I**). Notably, the 2 previous 2TEL-T4-lysozyme crystals also featured triangular connections brought about by 6 copies of the lysozyme packing together within a ring of 6 x 2TEL polymers (**Figure 1D,E**)[6]. The previous 3TEL-FUR crystals may have been well-ordered because the FUR domain makes contacts to 3 other 3TEL polymers in addition to its host polymer (**Figure 1F**), a feature also seen in the 3TEL-rigid-DARPin structure (**Figure 4G,I**)[7].

We were surprised to discover that the TELSAM polymers themselves need not directly contact one another in the crystal lattice. This is a profound discovery because removal of the requirement for adjacent TELSAM polymers to make direct contact should dramatically extend the upper size limit of target proteins that can be crystallized using TELSAM and supports continued use of the 1TEL (6 copies of the target protein per helical turn) construct. We have also shown that TELSAM-target protein fusions can form well-ordered, diffracting crystals using either flexible or semi-rigid TELSAM-target linkers. Future work is needed to determine whether each of 1TEL and 3TEL work better with rigid or flexible linkers, whether there is a complex relationship between the optimal number of target proteins displayed per helical turn and the linker type, and whether the optimal combination is specific for each target protein. The ability to use flexible linkers is compelling because it enables the possibility of designing TELSAM–target fusions based only on the predicted secondary structures of target proteins for which the tertiary structure is unknown. Future work is needed to define the maximum flexible linker length that reliably allows the formation of well-ordered crystals. The possibility of using longer TELSAM–target protein linkers is supported by our observation that the target protein can find a low energy, rigid binding mode against its own TELSAM polymer or an adjacent TELSAM polymer.

We have provided further evidence that the helical rise of the TELSAM polymer can adjust to allow target protein crystal contacts sufficient to form diffracting crystals and report a new minimum helical rise for a single-helical polymer of 46.9 Å. Further work is needed to determine whether rigidifying the helical rise of TELSAM is enabling or detrimental to the formation of ordered crystals and whether this is target protein-dependent.

In the study by Nauli, et al., the authors reported that many of the tested 2TEL–target protein fusions gave crystals with high mosaicity [6]. We did not observe mosaicity or other disorder issues in 1TEL-flex-vWa crystals, which also had very low anisotropy in their diffraction limits and for which 67% of the reflections could be indexed into the P65 space group. Conversely, the 3TEL-rigid-DARPin exhibited smeared reflections at certain diffraction angles (**Figure 2L**) and a high degree of anisotropy in the diffraction limits along each unit cell axis. In addition, only 12% of reflections could be indexed into the P1211 space group, suggesting a significant number of alternative lattices and disorder in this crystal. This disorder could be responsible for the extremely thin plate morphology of these 3TEL-rigid-DARPin crystals. The disorder may be due to the specific target protein, valency, linker type, or specific crystal lattice obtained, although DARPins are known to give well-ordered crystals, reasonably ruling out the target protein in this case.

Thus far, the TELSAM-mediated structures we have reported did not arise from diffraction data extending beyond 2 Å. Previously reported crystal structures of TELSAM–target protein fusions likewise have never exhibited resolution beyond 2.30 Å [6, 7]. Further work is needed to determine whether fusion to TELSAM imposes a limit on the resolution of resulting diffraction datasets and whether that limit can be extended through modifications to the TELSAM–target connection. We note that synchrotron radiation (SSRL beamline 9-2), a Pilatus 6M pixel array detector (Dectris), and the Autoproc software [10] were instrumental in mitigating the disorder in the 3TEL-rigid-DARPin crystals and extending the resolution of each of the structures reported here.

## Acknowledgments

Use of the Stanford Synchrotron Radiation Lightsource, SLAC National Accelerator Laboratory, is supported by the U.S. Department of Energy, Office of Science, Office of Basic Energy Sciences under Contract No. DE-AC02-76SF00515. The SSRL Structural Molecular Biology Program is supported by the DOE Office of Biological and Environmental Research, and by the National Institutes of Health, National Institute of General Medical Sciences (including P41GM103393). The contents of this publication are solely the responsibility of the authors and do not necessarily represent the official views of NIGMS or NIH.

